# Cooperativity between the 3’ untranslated region microRNA binding sites is critical for the virulence of eastern equine encephalitis virus

**DOI:** 10.1101/649525

**Authors:** Derek W. Trobaugh, Chengqun Sun, Nishank Bhalla, Christina L. Gardner, Matthew Dunn, William B. Klimstra

## Abstract

Eastern equine encephalitis virus (EEEV), a mosquito-borne RNA virus, is one of the most acutely virulent viruses endemic to the Americas, causing between 30% and 70% mortality in symptomatic human cases. A major factor in the virulence of EEEV is the presence of four binding sites for the hematopoietic cell-specific microRNA, miR-142-3p, in the 3’ untranslated region (3’ UTR) of the virus. Three of the sites are “canonical” with all 8 seed sequence residues complimentary to miR-142-3p while one is “non-canonical” and has a seed sequence mismatch. Interaction of the EEEV genome with miR-142-3p limits virus replication in myeloid cells and suppresses the systemic innate immune response, greatly exacerbating EEEV neurovirulence. The presence of the miRNA binding sequences is also required for efficient EEEV replication in mosquitoes and, therefore, essential for transmission of the virus. In the current studies, we have examined the role of each binding site by point mutagenesis of seed sequences in all combinations of sites followed by infection of mammalian myeloid cells, mosquito cells and mice. The resulting data indicate that both canonical and non-canonical sites contribute to cell infection and animal virulence, however, surprisingly, all sites are rapidly deleted from EEEV genomes shortly after infection of myeloid cells or mice. Finally, we show that the virulence of a related encephalitis virus, western equine encephalitis virus, is also dependent upon miR-142-3p binding sites.

**Author Summary:** Eastern equine encephalitis virus (EEEV) is one of the most acutely virulent mosquito-borne viruses in the Americas. A major determinant of EEEV virulence is a mammalian microRNA (miRNA) that is primarily expressed in myeloid cells, miR-142-3p. Like miRNA suppression of host mRNA, miR-142-3p binds to the 3’ untranslated region (UTR) of the EEEV genome only in myeloid cells suppressing virus replication and the induction of the innate immune response. In this study, we used point mutations in all four miR-142-3p binding sites in the EEEV 3’ UTR to understand the mechanism behind this miRNA suppression. We observed that decreasing the number of miR-142-3p binding sites leads to virus escape and ultimately attenuation *in vivo*. Furthermore, another virus, western equine encephalitis virus, also encodes miR-142-3p binding sites that contribute to virulence *in vivo*. These results provide insight into the mechanism of how cell-specific miRNAs can mediate suppression of virus replication.

## Introduction

Eastern equine encephalitis virus (EEEV) is a mosquito-borne alphavirus that causes severe manifestations of encephalitis in humans resulting in high mortality rates and long-term neurological sequelae in symptomatic cases [1] and is one of, if not the most acutely virulent virus circulating in North America. Even though both EEEV and the closely related Venezuelan equine encephalitis virus (VEEV) cause encephalitic disease, they exhibit drastic differences in their pathogenesis, in large part due to differential infectivity of the viruses for myeloid cells. VEEV is highly myeloid cell tropic and induces a robust innate immune response while EEEV is largely replication defective in these cells, which limits the production of systemic IFN-α/β and other innate cytokines *in vivo* [2]. Critical to EEEV virulence, a host microRNA (miRNA), miR-142-3p, primarily expressed in hematopoietic cells [3], prevents translation of the EEEV genome and virus replication in myeloid cells by interaction with binding sites in the EEEV 3’ untranslated region (UTR) [4]. Remarkably, although mosquitoes do not express miR-142-3p, presence of the miR-142-3p binding site sequences is required for efficient mosquito infection and, presumably, transmission of EEEV in nature [4].

miRNAs are short ~22 nucleotide long non-coding RNAs that can be transcribed from both coding and non-coding chromosomal regions. miRNAs bind to complimentary sequences in host mRNA as part of the RNA-induced silencing complex (RISC) through the seed sequence, nucleotides 2-8 at the 5’ end of the miRNA [5]. Perfect complementarity throughout the entire miRNA and target mRNA can lead to degradation of the mRNA, but is rarely seen in mammalian interactions [6,7]. More commonplace in mammalian cells is a perfect match (canonical sequence) or a single mis-match (non-canonical sequence) in the seed sequence and the target mRNA [8,9] leading to translational repression, downregulation of protein expression, and RNA destabilization [10,11]. The miR-142-3p blockade virtually eliminates EEEV translation and replication in myeloid cells [4].

miRNAs can interact with the 5’ UTR (e.g. hepatitis C virus (HCV) [12]), 3’ UTR (e.g. EEEV [4], and bovine viral diarrhea virus (BVDV) [13]) or coding regions (e.g. influenza [14–16], and enterovirus-71 [17,18]) (reviewed in [19]). This interaction leads to either inhibition of virus translation and replication (e.g. EEEV [4]) or stabilization of the virus RNA and increased replication (e.g. HCV [20] and BVDV [13]). There are three canonical and one non-canonical miR-142-3p sites in a 260 nt stretch of the EEEV 3’ UTR [4]; however, the contribution of each site to replication inhibition is not known, and the contribution of individual or combinations of miRNA binding sites to virulence in animals is not known for any virus.

In the current studies, we have leveraged the highly restrictive effect of miR-142-3p on EEEV replication *in vitro* and *in vivo* to examine the impact of each of the four sites present in the 3’ UTR on virus replication and disease. We have used point mutations to alter each of the miRNA binding sites to a non-functional sequence, either individually or in combination. We then examined infection efficiency for myeloid cells or mouse lymphoid tissue, replication, innate immune response induction, disease severity after mouse infection and competence for mosquito cell replication. We found that both canonical and non-canonical sites contributed to the virulence phenotype of wild type (WT) EEEV with a single site exerting the most potent effect upon virulence. The contribution of all sites to restriction of myeloid cell replication was reinforced by the observation that deletion mutations in the 3’ UTR arose rapidly during WT EEEV replication *in vitro* and in mice removing all of the miR-142-3p sites and conferring myeloid cell tropism to EEEV. Finally, we provide evidence that another encephalitic alphavirus, western equine encephalitis virus (WEEV) also possesses functional miR-142-3p binding sites in its 3’ UTR.

## RESULTS

### miR-142-3p binds to the EEEV 3’ UTR within the RISC complex in RAW cells

The RNA induced silencing complex (RISC) facilitates repression of host mRNA and viral RNAs through interactions of Argonaute protein (Ago) with host miRNAs [20,21]. In an effort to determine if miR-142-3p bound to viral RNA and targeted it to the RISC complex, we utilized a translation reporter that encodes the 5’ UTR and the WT EEEV 3’ UTR encoding the four miR-142-3p binding sites (Figure 1A) or the 11337 deletion mutant 3’ UTR that lacks all of the miR-142-3p binding sites [4] and two different experimental methodologies: 1) immunoprecipitation of the Ago protein to isolate RNAs interacting with the complex and 2) precipitation of biotin-labeled miR-142-3p to ascertain whether miR-142-3p interacted directly with the EEEV 3’ UTR in myeloid cells. Virus-specific RT-PCR of precipitated nucleic acids was then used to quantify the level of reporter RNA.

**Figure 1:**
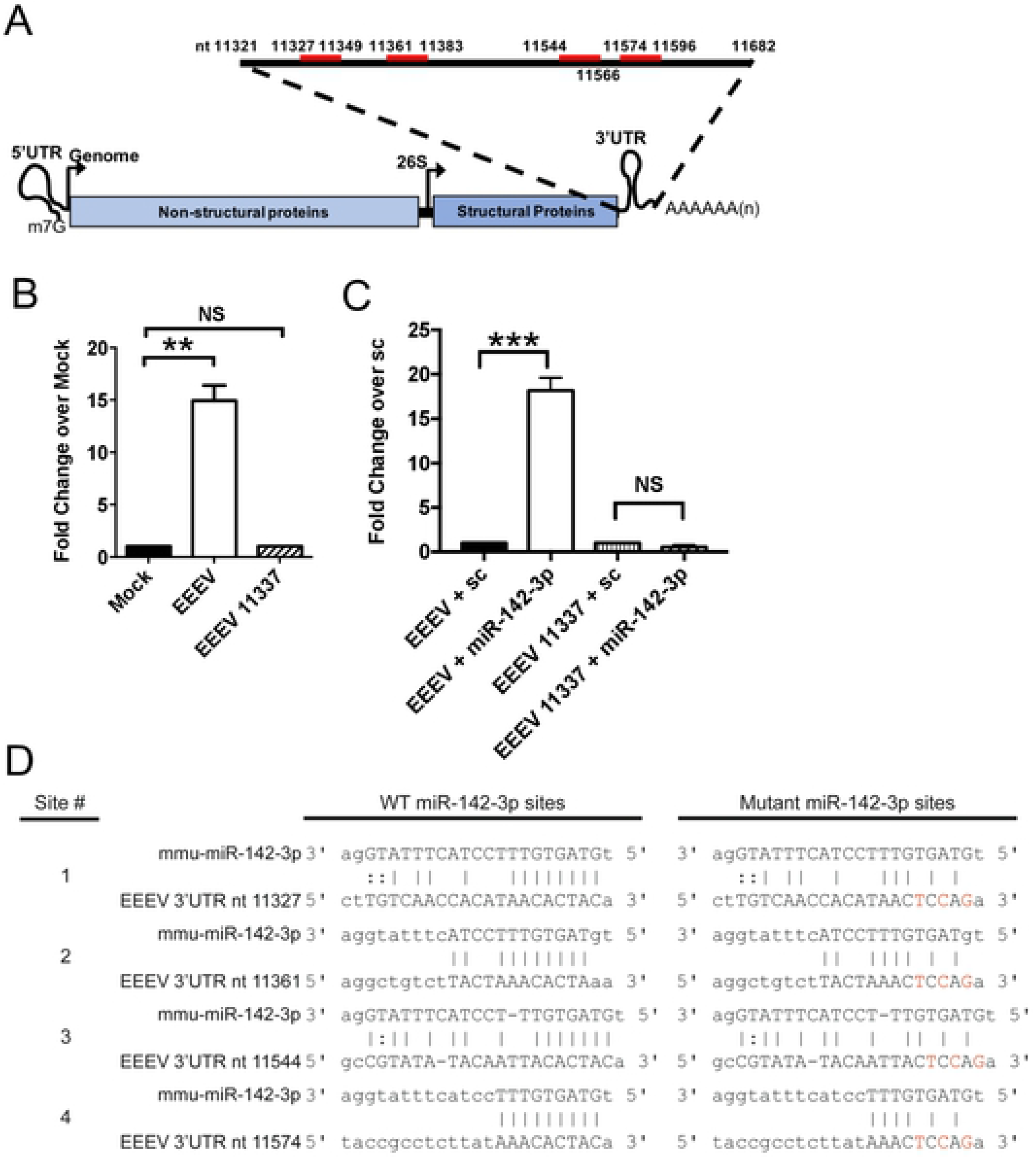
miR-142-3p binds to EEEV 3’ UTR in myeloid cells. A) EEEV genome structure. miR-142-3p binds (red bar) to EEEV genome in the 3’ UTR. Numbers indicate nucleotide position in the EEEV genome of the 3’ UTR and miR-142-3p binding sites B) EEEV or EEEV 11337 translation reporters (7ug) were electroporated into the RAW monocyte/macrophage cell line. Anti-Ago1/2 antibody was used to immunoprecipitation of Argonaute proteins. qRT-PCR was used to quantify amount of reporter RNA in each immunoprecipitated fraction. Data is represented as fold change over mock. **P<0.01, NS: not significant unpaired t test. n=2 independent experiments. Error bars represent SD. C) Biotin labeled miR-142-3p mimic or biotin labeled scramble (sc) were co-electroporated with the reporters into RAW cells. Streptavidin beads was used for immunoprecipitation and qRT-PCR was to quantify the amount of reporter RNA in each fraction. Data is represented as fold change over sc. n=2-3 independent experiments. Error bars represent SD. ***P<0.001, unpaired t test. D) Alignment of EEEV ‘3 UTR to mmu-miR-142-3p with red letter indicate mutated nucleotides (nt) in seed sequence. nt number indicates initial position in EEEV genome of miR-142-3p binding sites.

After electroporation of the EEEV translation reporters into RAW 264.7 monocyte/macrophage cell line, which constitutively expresses miR-142-3p[4], WT EEEV RNA was ~15 fold higher in the Ago-immunoprecipitated fraction compared to 11337 reporter RNA when normalized to mock electroporated cells and total input RNA (Figure 1B) demonstrating that WT EEEV interacted with the Ago complex. Next, to determine specifically if miR-142-3p bound to the EEEV 3’ UTR, we electroporated a biotin-labeled miR-142-3p or biotin-labeled scrambled control miRNA into RAW cells along with the EEEV reporters and immunoprecipitated RNA complexes with streptavidin beads. We detected ~18 fold higher levels of WT EEEV RNA in lysates that were co-transfected with a biotin-labeled miR-142-3p mimic compared to electroporation with a scrambled miRNA mimic (Figure 1C). Furthermore, we detected only very low levels of 11337 reporter RNA in lysates after co-transfection with both the biotin-miR-142-3p mimic or scrambled mimic demonstrating the specificity of miR-142-3p interaction with the WT EEEV 3’ UTR. PCR analysis indicated that the initial levels of WT and 11337 reporter RNA in electroporated RAW cells were comparable prior to immunoprecipitation. Thus, the EEEV 3’ UTR interacts with miR-142-3p inside RISC complex through the Ago proteins presumably leading to translational inhibition of the EEEV genome.

### Combinatorial mutation of the EEEV miR-142-3p binding sites increases virus replication in myeloid cells

A higher number of miRNA binding sites within a virus 3’ UTR may lead to increased suppression of tissue-specific virus replication and virus attenuation *in vivo* [22,23]. To determine the contribution of each miR14-2-4p binding site to restriction of EEEV replication, we created point mutant EEEV viruses in every combination of one, two, three or all four miR-142-3p binding sites disrupted in the 3’ UTR (Figure 1D) in the background of a nLuc expressing virus [24]. We incorporated mutations at 3 nucleotides at positions 2 (C2G), 4 (T4C), and 6 (A6T) in each of the canonical miR-142-3p binding sites that is complimentary to the miR-142-3p seed sequence (Figure 1D, sites 1, 3 and 4). We also made similar mutations in the non-canonical miR-142-3p binding site (site 2), except at position 2 where an A-G mutation was inserted making all four binding sites have the same 3 nucleotide mutations. This yielded fifteen mutant viruses with the combined four-site mutant containing a total of 12 point mutations over a span of ~250 nucleotides.

In BHK-21 fibroblast cells, which do not express miR-142-3p [4], point mutation of the miR-142-3p binding sites did not significantly change virus replication compared to WT EEEV or the 11337 deletion mutant at 12 hpi (Figure 2A) or subsequent time points (data not shown). In RAW cells, a monocyte/macrophage cell line which express an intermediate level of miR-142-3p, and C57BL/6 bone marrow-derived dendritic cells (BMDCs), which express a high level of miR-142-3p [4], mutation of each miR-142-3p binding site individually, leaving three intact miR-142-3p binding sites, did not significantly increase virus replication compared to WT EEEV (Figure 2B-C). Mutation of any two of the miR-142-3p binding sites resulted in a reproducible but small increase in virus replication in both RAW and BMDCs that did not attain statistical significance *versus* WT EEEV. However, mutation of 3 miR-142-3p binding sites, leaving only a single binding site intact, led to significantly higher levels of virus replication at 12 hpi in both RAW and BMDC cells (Figure 2B-C), with the exception of the virus with mutations in sites 1, 2, and 3 leaving site 4 intact in RAW cells.

**Figure 2:**
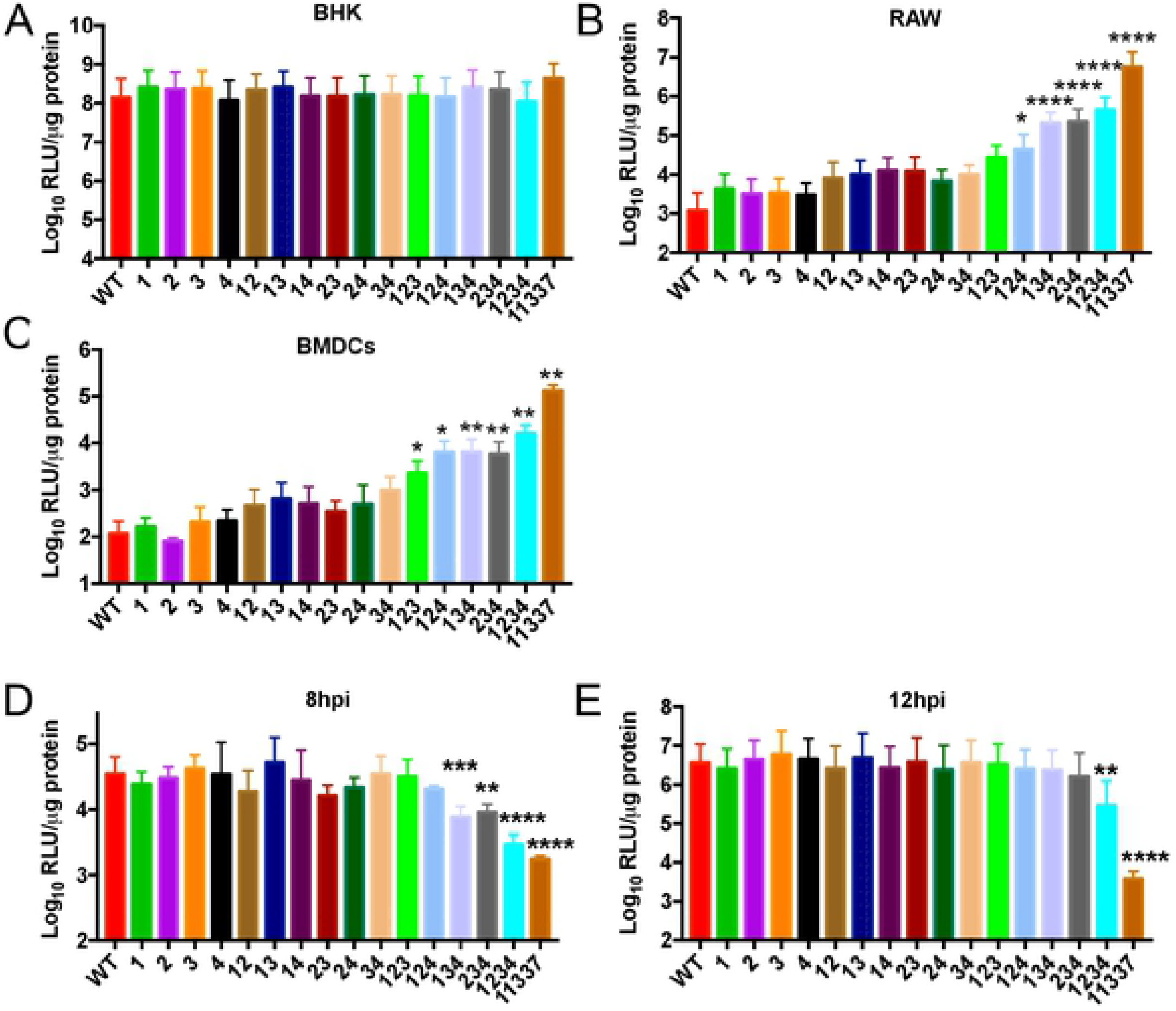
Decreasing the number of miR-142-3p binding sites leads to increased virus replication in myeloid cells and viral attenuation *in vivo.* A-C) Quantification of virus replication at 12 hpi. BHK (MOI=0.1), RAW (MOI=0.1), BMDCs (MOI=5) were infected with the EEEV mutants expressing nLuc. D-E) Virus replication in C6/36 mosquito cells (MOI=1) at (D) 8hpi and (E) 12hpi. Data is log_10_ transformed and expressed as relative light units (RLU) per μg protein. N=6-9, 2-3 independent experiments. *P<0.5, **P<0.01, ***P<0.001, ****P<0.0001 one way ANOVA with corrections for multiple comparisons using the Holm-Sidak method comparing each mutant to WT. Error bars represent SE.

Both the 11337 deletion mutant and the 1234 point mutant are impaired in their ability to replicate in *in vitro* in C6/36 mosquito cells and in mosquito vectors [4], suggesting a requirement for these 3’UTR sequences in the EEEV transmission cycle. Therefore, we used the point mutant viruses to determine whether or not the individual miR-142-3p binding sites contribute to EEEV replication in mosquito cells. At 8 hpi, replication of both the quadruple mutant 1234 and 11337 mutant viruses were significantly reduced in C6/36 cells compared to EEEV WT (Figure 2D). Of the three mutation viruses, only mutants 134 and 234 were significantly reduced compared to EEEV WT, while there was no difference in replication of the other EEEV mutants. By 12hpi, only mutants 1234 and 11337 were significantly reduced compared to WT (Figure 2E). Combined with the fact that the 11337 mutant was significantly more inhibited than the four site point mutant at 12 hpi (p<0.0001, unpaired t test), this result suggests that the individual miR-142-3p binding site sequences may not be the primary elements in the 3’ UTR that promote mosquito replication, but potentially, a sequence-dependent RNA secondary structure or other specific sequence is critical.

Together, these data demonstrate that while mutations in the miR-142-3p binding sites do not alter virus replication in non-miR-142-3p expressing cells, mutations in 3 of the miR-142-3p binding sites results in EEEV replicative escape from miR-142-3p suppression in myeloid cells. Therefore, 2 or more miR-142-3p binding sites are required for significant suppression of EEEV myeloid cell replication *in vitro* demonstrating that cooperativity between the miR-142-3p binding sites enhances EEEV suppression.

### Combinatorial mutation of the EEEV miR-142-3p binding sites increases virus attenuation in mice

We have previously demonstrated that deletion of all of the miR-142-3p binding sites (virus 11337) resulted in virus attenuation *in vivo* due to increased myeloid cell replication and systemic IFN-α/β production [4]. Since decreasing the number of functional miR-142-3p binding sites resulted in increased myeloid cell replication *in vitro,* we sought to determine whether a similar phenomenon could be observed *in vivo*. In CD-1 mice infected with viruses lacking a single miR-142-3p binding site (single mutants), only mutation of site 4 lead to a significant increase in survival compared to WT EEEV (Figure 3A); there were no significant survival differences with mutation of sites 1, 2, or 3 alone. Mutation of two miR-142-3p binding sites (double mutants) resulted in greater attenuation and higher percent survival compared to the single mutants; however, virus attenuation was dependent on the combination of sites that were mutated. The virus with sites 3 and 4 mutated (34) had the highest survival rate of the double mutants and was significantly attenuated with 50% survival compared to WT EEEV (Figure 3A). Percent survival was also significantly higher in mice infected with the 13, 14, and 23 viruses compared to WT EEEV but lower than the 34 mutant. There was no significant survival difference in mice infected with the 12 and 24 mutant viruses compared to WT EEEV.

**Figure 3:**
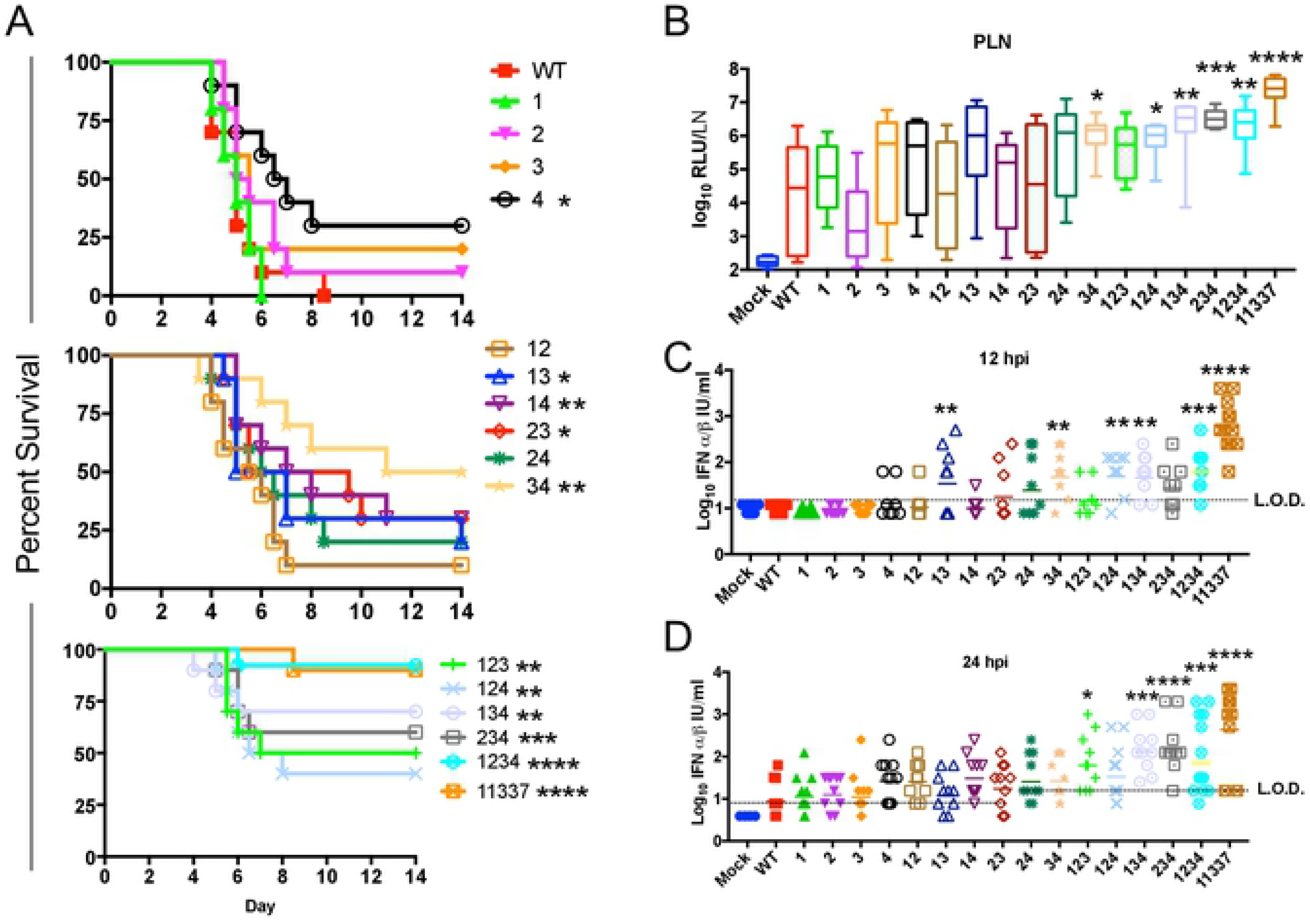
Elimination of miR-142-3p binding sites leads to attenuation due to increased virus replication in popliteal lymph node and serum IFN. A) Female outbred CD-1 mice (5-6 weeks) were infected with 10^3^ pfu sc in each footpad. Morbidity and mortality were measured twice daily. n=10 mice from 2 independent experiments *P<0.5, **P<0.01, **P<0.001, ****P<0.0001, Log-Rank Test comparing each EEEV mutant to WT. B) Outbred CD-1 mice were infected with 10^3^ pfu of WT, 11337 or each mutant sc in each footpad. A single popliteal lymph node (PLN) was harvested at 12 hpi and analyzed for nLuc expression. Data is log_10_ transformed and represented as RLU per LN. n = 7-12 mice, from 2-3 experiments. Box-and whisker plots represent min-max with bar representing the median value. C-D) Biologically active IFN-α/β levels in serum at 12 hpi (B) and 24 hpi (C). CD-1 mice were infected with 10^3^ pfu of WT, 11337 or each mutant sc in each footpad. n=8-12 mice, from 2-3 experiments. *P<0.05, **P<0.01, ***P<0.001, ***P<0.0001, comparing the log-transformed data of each mutant with WT using the one way analysis of variance test with corrections for multiple comparisons using the Holm-Sidak method. LOD = limit of detection of the IFN assay. Bar represents geometric mean.

Infection of mice with viruses encoding three mutant miR-142-3p binding sites and only one functional miR-142-3p binding site (triple mutants) resulted in significant survival differences for all of the mutants compared to WT EEEV (Figure 3A). Additionally, percent survival for the mutant viruses 134 and 234 was higher than that of the double mutants, but not to the level of 11337 or the quadruple mutant 1234 virus. Finally, mice infected with the mutant 1234 virus was not significantly different than 11337. We also observed similar survival percentages of the miR-142-3p binding site mutants in inbred C57BL/6 mice compared to CD-1 mice (S1 Figure).

Together these results demonstrate that, similar to our data on virus replication in myeloid cells, reduction in the number of miR-142-3p binding sites leads to increased survival and higher virus attenuation *in vivo*. Importantly, none of the 1, 2, or 3 site point mutants were similarly attenuated to the four-site mutant 1234 or 11337, suggesting each of the four miR-142-3p binding sites, including the non-canonical site (site 2) contributes to the fully virulent phenotype of the WT virus. There was little significant difference in attenuation between viruses with similar numbers of mutations. Mutant virus 4 was significantly attenuated compared to mutant 1 (P<0.05) and mutant 34 virus was significantly attenuated compared to 12 (P<0.01), which may suggest a higher impact of site 4 that other sites on virulence in vivo. Yet overall, the data indicate that each miR142-3p binding site contributes to EEEV virulence.

### Early virus replication in popliteal lymph nodes and inflammatory response is dependent on the number of miR-142-3p binding sites

We also previously demonstrated that the miR-142-3p binding site mutant, 11337, rescued virus replication in the popliteal lymph nodes (PLN) *in vivo* early after infection compared to WT EEEV [4]. Therefore, we next determined whether or not combinatorial mutation of the miR-142-3p sites led to differences in virus replication in the PLNs early after infection. CD-1 mice were infected with the nLuc-expressing EEEV mutant viruses and the PLNs were harvested 12 hpi, and virus replication was quantified by nLuc analysis. In the PLNs, the deletion mutant 11337 virus had the highest level of virus replication at 12 hpi (Figure 3B), which was significantly higher than WT. Replication of the four site mutation virus, 1234, and the triple mutant viruses were higher than either the single or double mutant viruses, but lower than the 11337 mutant virus. All of the triple mutant viruses except mutant 123 had significantly higher levels of virus replication in the PLN compared to WT although that mutant exhibited a trend toward higher replication. Of the double mutants, only mutants 34 (P<0.05) had significantly higher replication than WT. Replication of the single site mutation viruses was not significantly different form the WT, although the site 4 mutant exhibited a trend towards higher replication evidenced in some animals. These results are similar to the *in vitro* results in myeloid cells in that the mutant viruses with 2 or 3 functional miR-142-3p binding sites showed restricted viral replication. In summary, elimination of 3 miR-142-3p binding sites conferred myeloid cell replication leading to higher virus levels of nLuc in the PLN. Virus replication *in vivo* was more variable after infection than in cultured myeloid cells suggesting that other factors potentially including variability in the expression of miR-142-3p in PLN cells or numbers of myeloid cells could be influencing PLN replication.

Myeloid cell replication by the mutant 11337 virus leads to IFN-α/β production *in vivo* by 12 hpi [4]. We hypothesized that decreasing the number of miR-142-3p binding sites would lead to higher levels of systemic IFN-α/β contributing to the virus attenuation that is seen *in vivo*. At 12 hpi, serum IFN-α/β, measured by bioassay, was undetectable in WT and single mutation viruses with the exception of two mice infected with the site 4 mutant (Figure 3C). This is consistent with the trend for site 4 mutant viruses towards higher PLN replication and significant attenuation *in vivo* demonstrating that activity of 3 miR-142-3p binding sites largely suppresses IFN-α/β production except when the miR-142-3p binding site (site 4) is closest to the poly (A) tail. Serum IFN-α/β levels in mice infected with the double mutant viruses was dependent on the particular combination of mutations. Mutant viruses 12 and 14 only had a single mouse with detectable levels of IFN-α/β. Viruses with mutations in sites 23, and 24 led to a higher number of mice with serum IFN-α/β, but the average for the groups was not significantly different from WT. Mutant 13 and 34 were the only double mutant viruses to have significantly higher levels of serum IFN-α/β compared to WT (P<0.01). For the triple mutants, mutant 123 and 234 didn’t not induce significantly higher levels of IFN-α/β compared to WT.

By 24 hpi (Figure 3D), we detected serum IFN-α/β in some mice infected with the WT, and more mice infected with the single mutants and the double mutants than at 12hpi. These data suggest there is a delayed induction of IFN-α/β with these viruses compared to 11337 and the other mutants. It is possible that the increased IFN-α/β production by WT may be due to changes in miR-142-3p levels within the PLN as a response to infection, leading to virus replication or, possibly, virus escape of miR-142-3p suppression in myeloid cells due to the presence of higher levels of WT virus in the PLN at 24 hpi compared to 12 hpi [25].

Early production of cytokines and chemokines produced by myeloid cells can influence trafficking of immune cells and the induction of the adaptive immune response [26,27]. A single PLN was harvested at 12 hpi from each of three CD-1 mice infected with either WT, 11337 or mock-infected and RNA was isolated for qRT-PCR to quantify cytokine and chemokine mRNA levels using a panel of cytokine and chemokines that are involved in trafficking and induction of the adaptive immune response, specifically trafficking of immune cells to peripheral tissues during infection [26]. Significantly higher mRNA levels of the inflammatory cytokines *Ifnb, Ifng, Il6, and Il1b* and chemokines *Cxcl10*, *Ccl3,* and *Ccl2* were detected in 11337-infected PLN compared to WT-infected PLN. (S2A Figure). For both *Ccl4* and *Cxcl1*, statistical significance was influenced by several WT-infected mice that had high mRNA levels in the PLN, but overall WT-infected mice had lower levels compared to 11337-infected mice.

To further examine the relationship of myeloid cell replication and the number of miR-142-3p binding sites to induction of chemokine mRNAs within the PLN, we quantified the level of *Cxcl10* mRNA within the PLN after infection with each of EEEV mutants at 12 hpi (S2B Figure). Similar to IFN-α/β levels in the serum, *Cxcl10* mRNA levels in the PLN depended on the number of miR-142-3p binding sites. Mice infected with WT and the single mutants had similar levels of *Cxcl10* mRNA within their PLN. Mice infected with the double mutant virus 34 had significantly (P<0.001) higher levels of *Cxcl10* mRNA compared to WT. Mutant 123 of the triple mutant viruses didn’t not significantly increase *Cxcl10* mRNA within the PLN, but all of the other triple mutant viruses, the quadruple mutant 1234, and 11337 had significantly higher levels of *Cxcl10* mRNA. Together, our results demonstrate the increasing myeloid cell replication leads to not only higher levels of serum IFN-α/β, but also higher mRNAs levels of inflammatory cytokines and chemokines in the PLN. Overall, these results demonstrate that, similar to myeloid cells *in vitro*, the four miR-142-3p binding sites in the EEEV 3’ UTR have a cumulative effect upon PLN replication and response, and overall virulence, However, deficiency in site 4 does have a consistent but not always significantly greater effect upon these factors.

### Virus-cell interaction in the PLN influences replication in CNS

As we have demonstrated above, reducing the number of miR-142-3p binding sites in the EEEV 3’ UTR leads to increased virus attenuation *in vivo*. An important question regarding the pathogenesis of arboviral encephalitic viruses is whether or not attenuation of encephalitis-causing viruses can be influenced by responses elicited in tissues outside the CNS. This is particularly significant with EEEV as we have proposed that failure to elicit a robust peripheral innate immune response contributes to the extreme virulence of the virus. Tissues from CD-1 mice infected with the EEEV mutants were harvested 96 hours post infection to measure virus replication in the PLN, spleen, and several regions of the brain. In the PLN and spleen, the triple, quadruple and 11337 mutant viruses all had lower levels of mean virus replication compared to WT (S3 Figure). These mutants also elicited higher levels of serum IFN-α/β than WT at 12 hpi. There was no difference in virus replication between WT and the single or double mutants at this time point suggesting that early IFN-α/β production by the triple, quadruple, and 11337 (Figure 3C), in the absence of complete miR-142-3p restriction, led to reduced virus replication *in situ* in the PLN and spleen.

We previously demonstrated that higher levels of serum IFN-α/β lead to increased STAT1 phosphorylation and, presumably, upregulation of the IFN-induced antiviral state in brain tissue at early times post EEEV infection prior to neuroinvasion by the virus [28]. At 96 hours post infection with the EEEV mutants, we harvested the cortex, subcortex, cerebellum and olfactory bulbs to measure virus replication by nLuc analysis. WT-infected mice exhibited high levels of virus replication in all brain regions (Figure 4A-D) corresponding to the time when WT EEEV infected mice begin to succumb to infection (Figure 3A). Similar levels of virus replication were also detected in the different regions of the brain infected with the single mutants and double mutants. Even though all of the mice infected with the double mutant viruses had virus replication in the different regions of the brain, not all mice succumbed to infection (Figure 3A).

**Figure 4:**
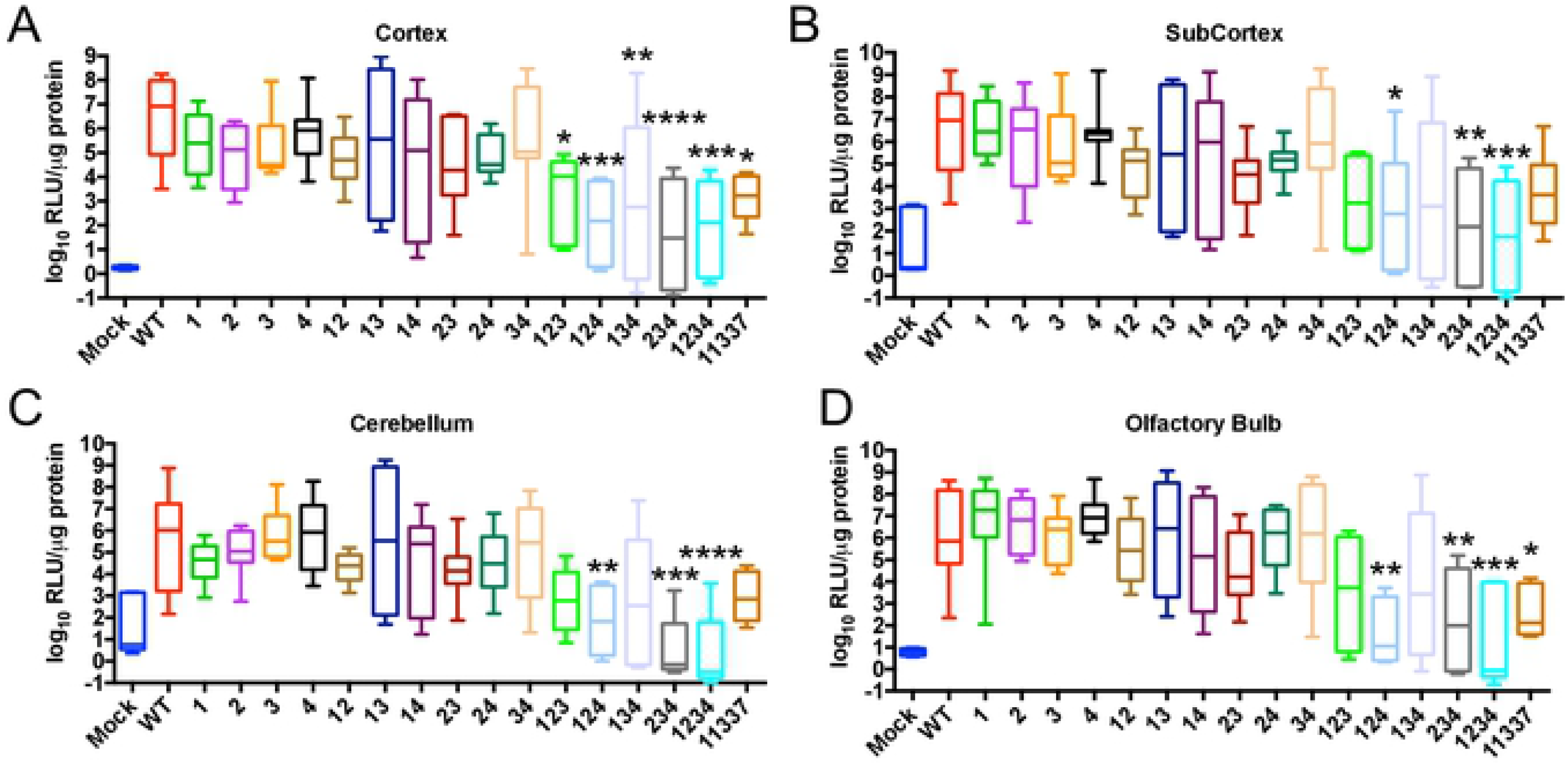
Reduced virus replication in the central nervous system after infection with the triple and quadruple mutant EEEV viruses. CD-1 mice were infected with 10^3^ pfu of the EEEV mutants sc in each footpad. Tissues were harvested at 96 hours post infection. Virus replication in cortex (A), subcortex (B), cerebellum (C), and olfactory bulb (D). Data is log_10_ transformed and represented as either RLU/LN or RLU per μg protein N=8 mice, from 2 independent experiments. *P<0.5, **P<0.01, ***P<0.001, ****P<0.0001 one way analysis of variance test with corrections for multiple comparisons using the Holm-Sidak method comparing each mutant to WT. Box-and whisker plots represent min-max with bar representing the median value.

Virus replication was significantly lower for the triple mutants, the quadruple mutant 1234, and the deletion mutant 11337, compared to WT in all four regions of the brain. In fact, virus replication of these mutants was also lower than the single and double mutants at this time point. As further evidence of the contribution of extraneural replication to the attenuated phenotype of binding site mutants, we found that the WT and the most attenuated 11337 mutant viruses caused similar virulence when give intracerebrally to CD-1 mice (S4 Figure). These results demonstrate that the miR-142-3p binding sites not only suppress virus replication in the PLN early after infection, but the IFN-α/β induced early after infection reduces virus dissemination to the CNS contributing to the attenuation of these viruses *in vivo*.

### Spontaneous mutation miR-142-3p binding sites may contribute to changing EEEV phenotypes as infection progresses

EEEV is primarily transmitted between ornithophilic mosquitoes and passerine birds, but infections do occur in humans and horses, both of which are considered dead-end hosts [29] and, therefore, should not contribute to the host-driven evolution of the virus. Mosquito cell lines do not express miR-142-3p [4,30] while avian hematopoietic cell lines do [31]; however, it may be expressed at lower levels than in mammal hematopoietic cell lines [4]. Importantly, without the miR-142-3p binding sites in the 3’ UTR, EEEV cannot establish a productive infection in mosquitoes [4], suggesting that the miR-142-3p binding sites are maintained due to selective pressures in the mosquito-bird life cycle but not during replication in humans or possibly mice as human disease models. Remarkably, the sequence of the region of the EEEV 3’UTR containing the miR-142-3p binding sites is identical between currently circulating viruses and viruses isolated in the 1930’s, also suggesting a strong selective pressure for conservation [4]. Therefore, we sought to determine if some of the altering phenotype of EEEV during mouse infection, such as the increase in serum IFN-αβ levels in mice infected with the WT or single mutation viruses between 12 and 24 hpi (Figure 3C-D), might be associated with a reduction in miR-142-3p restriction in myeloid cell infection. Interestingly, in RAW cells and interferon non-responsive mouse BMDCs, between 24 and 48 hpi, WT EEEV appears to escape miR-142-3p suppression and replicates to high titers (Figure 5A-B).

**Figure 5:**
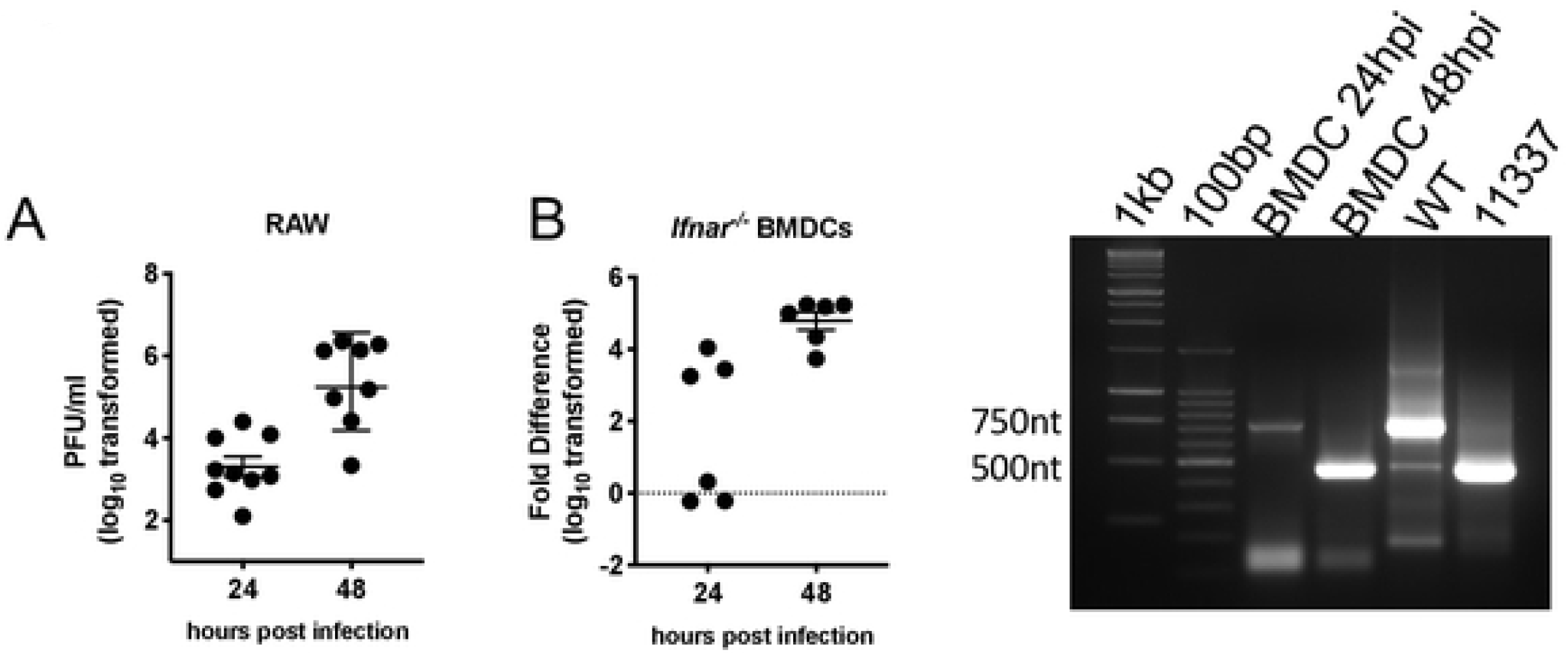
Escape of miR-142-3p suppression in myeloid cells. A-B) Replication of EEEV increases between 24 and 48 hpi *in vitro* detected by plaque assay in A) RAW cells or semi-quantitative RT-PCR in (B) *Ifnar*^−/−^ BMDCs. n=6-9 individuals wells from 2-3 independent experiments. Fold difference in B is compared to mock samples. Bar represents geometric mean and geometric SD. C) Representative gel from PCR of EEEV 3’ UTR of BMDC escape mutant between 24 hpi and 48 hpi. Control PCR product is approximately 750nt for WT EEEV and 500nt for 11337. 1kb and 100bp indicate 1kb and 100bp ladders respectively.

To detect escape mutations *in vitro* and *in vivo*, RNA was isolated from WT EEEV-infected RAW cells (48 hpi), *Ifnar*^−/−^ bone marrow derived dendritic cells (BMDCs) (24 hpi and 48 hpi), mouse brains, sera, and cervical lymph nodes (CVLN) at times >24 hours post infection. PCR amplification of the 3’ UTR led to the identification of a subset of amplified fragments in all amplified RNA that were smaller than the WT amplicon; however, with mouse samples, these fragments did not represent the majority of the viral population (data not shown). By comparison, in BMDCs, at 24 hpi a majority of the population was similar to WT (~750nt), but by 48 hpi, the predominant viral population had a much shorter 3’ UTR more similar to 11337 (Figure 5C). In general, 3’UTR length variations were of multiple lengths (S5 Figure). However, amplicon sequencing revealed that in all circumstances, deletions encompassed most or all four of the miR-142-3p binding sites and the sequence of one sample from the serum of an infected mouse was identical to the 11337 mutation (mutation 11337-11596). Two samples from the brains of B6 mice had identical deletions (11355-11601). These results demonstrate that the miR-142-3p binding sites are not stable during EEEV infection in a model of human disease.

### miR-142-3p binding sites are present in the EEEV-derived 3’ UTR of western equine encephalitis virus

Western equine encephalitis virus (WEEV) is a recombinant alphavirus consisting of the 5’ UTR, nonstructural proteins, capsid protein, and part of the 3’ UTR of EEEV and the structural proteins (E3, E2, and E1) and the initial 60 nucleotides (nt) of the SINV 3’ UTR [32,33]. Like EEEV, replication of WEEV is inhibited in human peripheral blood leukocytes[34]. Since part of the WEEV 3’ UTR is derived from an EEEV progenitor, we hypothesized that the 3’ UTR may also contain miR-142-3p binding sites. Screening of the WEEV 3’ UTR of the McMillan strain (McM) identified four potential miR-142-3p binding sites located at different positions relative to the poly (A) tract and each other (Figure 6A-B) as compared to the miR-142-3p binding sites in the EEEV 3’ UTR (Figure 1A). Two of the miR-142-3p binding sites, beginning at nt 11347 and 11369, have canonical seed sequence matches, but they overlap in the 3’ UTR by 6 nts. The other 2 miR-142-3p binding sites, beginning at 11405 and 11429, have a G:U wobble at position 2 of the seed sequence suggesting they may be non-canonical (Figure 6B).

**Figure 6:**
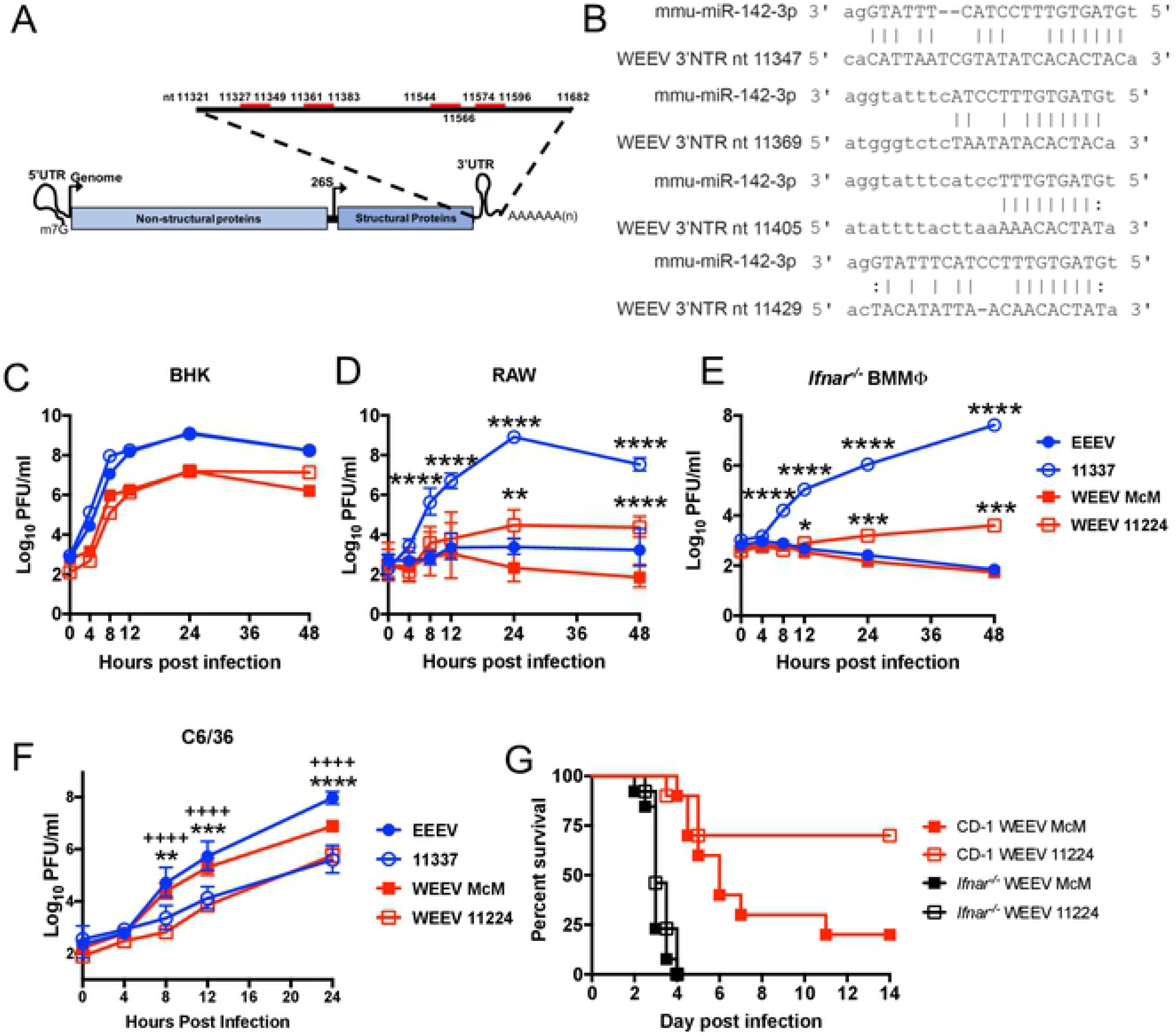
WEEV McMillan (McM) encodes miR-142-3p binding sites that restrict myeloid cell replication. A) Alignment in genome and location of miR-142-3p binding sites in WEEV McM 3’ UTR. B) Alignments between murine mmu-mIR-142-3p and WEEV 3’UTR. C-F) Viral growth curves in BHK (C), RAW (D), *Ifnar*^−/−^ BMMΦ (E), or C6/36 (F) of wild-type viruses (filled) or miR-142-3p mutants (open). Multiplicities of infection: BHK and RAW MOI=1, *Ifnar*^−/−^ BMMΦ MOI=5. n=6 from 2 independent experiments. Data was log_10_ transformed and and multiple unpaired *t*-tests were performed and corrected for multiple comparisons using the Holm-Sidak method. *p<0.05, **P<0.01, ***P<0.001, ****P<0.0001. ++++P<0.0001 comparing WEEV McM versus WEEV 11224 in C6-36 cells. Data is represented as geometric mean with geometric SD. G) Infection of outbred CD-1 mice (n=10 mice, 2 independent experiments) or *Ifnar*^−/−^ mice (n=13 mice, 2 independent experiments) with 10^4^ pfu of wild-type WEEV McM or mutant WEEV 11224 sc in both rear footpads. Morbidity and mortality were monitored twice daily. Mantel-Cox Log rank test was used to compare between viruses.

To determine whether the miR-142-3p binding sites within the WEEV 3’ UTR suppress WEEV replication in myeloid cells, we made a deletion mutant, WEEV-11224, that deletes 226 nucleotides from the 3’ UTR that eliminates all of the miR-142-3p binding sites similar to the 11337 deletion in EEEV[4]. In BHK cells, both WEEV McM and WEEV-11224 replicated with similar growth kinetics demonstrating that the deletion in the 3’ UTR had no effect on virus replication in miR-142-3p deficient cells (Figure 6C) [4,30]. In RAW cells that express miR-142-3p [4], both WEEV McM and WT EEEV did not replicate (Figure 6D). Deletion of the miR-142-3p binding sties in McM (WEEV 11224 virus) resulted in a 2-log increase in virus replication at 24 hpi compared to WEEV McM (P<0.01). By comparison, EEEV 11337 replicated to a ~4-log higher titer than WEEV 11224 at 24 hpi. Differences between EEEV and WEEV in relative growth of WT viruses and miR-142-3p deletion mutants may reflect differential sensitivity to mutant-induced interferon responses or other myeloid cell factors as similar responses were since in *Ifnar*^−/−^ BMMϕ (Figure 6E). Like EEEV 11337, WEEV 11224 is also inhibited in replicating in C6/36 mosquito cells (Figure 6F) suggesting a common function of this region between WEEV and EEEV in establishing mosquito replication. Finally, sc infection of mice revealed that the 11224 mutant was attenuated (P=0.052) *versus* the WT WEEV McM (25% versus 75% mortality, respectively) (Figure 6G). Attenuation of the EEEV miR-142-3p deletion mutant 11337 is due to differential type I IFN induction[4], therefore we infected *Ifnar*^−/−^ mice to determine whether WEEV 11224 was similarly attenuated by the type I IFN response. Survival times and mortality of *Ifnar*^−/−^ mice infected with WEEV McM and WEEV 11224 were not distinguishable (Figure 6G) demonstrating that type I IFN is needed for the attenuation of WEEV 11224 compared to WEEV McM. Thus, similar to EEEV, the miR-142-3p binding sites in the WEEV McM 3’ UTR restrict virus replication in myeloid cells leading to suppressed IFN responses and increased virulence in mice.

## DISCUSSION

miRNAs function in a cell and tissue specific manner to prevent translation of cellular mRNAs. Recent evidence demonstrates that cellular miRNAs can interact with RNA viruses to either suppress RNA translation via interaction with 3’ UTRs or enhance virus replication by interacting with 5’ UTRs (reviewed in [19]). Artificial insertion of miRNA binding sites into viral 3’ UTRs has been used to restrict virus replication in specific cells or tissues to understand the contribution of these cells and tissues to viral pathogenesis. These experiments require insertion of multiple highly complementary miRNA sequences in specific locations and distances from each other to achieve efficient viral suppression [22,23,30,35,36]. However, miRNA natural binding sites in viral 3’ UTRs are not completely complimentary nor are they arranged at uniform distances from each other [4]. We have used point mutations to determine the role of each of four naturally occurring canonical and non-canonical miR-142-3p sites whose stringent restriction of myeloid cell replication drives the severity of encephalitis caused by EEEV, one of the most acutely virulent viruses endemic to the Americas. Through these studies, we have confirmed that miR-142-3p interacts with the EEEV 3’ UTR as part of the RISC complex in myeloid cells *in vitro* and we have determined that all four of the miR142-3p sites, including the non-canonical site 2, contribute to EEEV virulence. Furthermore, we demonstrate that another encephalitic alphavirus, WEEV, achieves virulence through miR-142-3p restriction.

Viruses [22] and cellular mRNA [37–39] encoding multiple miRNA binding site are more suppressed than viruses and mRNA encoding only a single miRNA binding site. This interaction between two or more miRNA binding sites leading to enhanced repression of the target RNA over a single miRNA binding site is called cooperativity [38–40]. Cooperativity between miRNAs is most optimal when the seed sequences of the miRNA binding sites are spaced 13-35 nucleotides apart [40]. Also, the location of a miRNA binding site in the 3’ UTR within 15 nucleotides from the stop codon can increase efficacy and repression of the target RNA [38]. However, not all naturally encoded viral miRNA interaction domains encode more than one miRNA binding site [17,18,41,42], and if there is more than one miRNA binding site, the sites may not be ideally located based on the aforementioned guidelines for cooperativity.

For EEEV, the number of miR-142-3p binding sites rather than their location within the 3’ UTR was dominant in the suppression of EEEV replication. EEEV mutants with 2 or more miR-142-3p binding sites were suppressed in replication in myeloid cells *in* vitro to WT EEEV and *in vivo* in the PLN. Once a third miR-142-3p binding site was removed from the 3’ UTR leaving only one intact miR-142-3p binding site, EEEV virus replication was rescued in myeloid cells and the PLN. Even though replication was detected with the triple mutant viruses, it was still lower than the quadruple miR-142-3p mutant, 1234, and the deletion mutant, 11337 indicating that each miR-142-3p binding site contributes to EEEV restriction *in vitro*. However, it should be noted that mutant 4 exhibited a non-significant but consistent trend towards higher replication potentially implying a more substantial contribution to EEEV restriction that the other binding sites.

With *in vivo* morbidity/mortality studies, which appeared to distinguish more precisely between effects of individual binding sites, some of the single, double, and triple mutants were significantly attenuated in comparison with WT EEEV. Of the single mutants, the mutant EEEV virus lacking site 4 was the most attenuated. This miR-142-3p binding site is located the closest to the poly (A) tail, a known factor in enhancing miRNA restriction of cellular mRNA [38], and its position within the UTR may underlie greater replication restriction observed *in vitro* and *in vivo.* The mutant virus 34, which was mutated in two sites that are only separated by 8 nucleotides, was the most attenuated double mutant virus compared to WT EEEV. This suggests that there is some positional effect for these sites that may enhance cooperativity. Of the triple mutants, the viruses mutated in both sites 3 and 4 in conjunction with either site 1 or 2 were the most attenuated, again suggesting greater contribution of site 4 and an interaction between sites 3 and 4.

All of the triple mutants were attenuated, but more virulent than the quadruple mutant 1234 or 11337. Site two with a G:U wobble at position 2 of the seed sequence is a non-canonical miRNA binding site due to a potential offset 6-mer binding site that has complementarity between nucleotides 3-8 of the seed sequence and the EEEV 3’ UTR [43]. The nullification of this non-canonical site in the mutant 134 still resulted in increased mortality during infection compared to 1234 and 11337. Therefore, our data demonstrate that each miR-142-3p binding site, even the non-canonical site 2, is involved in suppression of virus replication in myeloid cells and attenuation of EEEV disease. However, as noted, site 4 may be the most restrictive by itself and positional effects between sites 3 and 4 may enhance cooperativity yielding the greater suppression of replication.

It can be argued that the binding of miRNAs to viral RNAs can lead to sequestration of the miRNA leading changes in the host transcriptome due to derepression of previously repressed targets [13,44] and with EEEV, potentially causing altered myeloid cell responses to infection. However, the binding of miRNAs to viral RNA is unlikely to result in changes in the cellular transcriptome through sequestration of the free miRNA early after infection when viral genomes are limited. We previously demonstrated with EEEV, that initial translation of the infecting virus genome is restricted by miR-142-3p and subsequent replication is greatly suppressed [4]. Here we show that EEEV RNA associates with Ago complexes and therefore, is directed into translation repression pathways. Furthermore if the miRNA is highly expressed within the cell, as with miR-142-3p in myeloid cells [3], the number of miR-142-3p molecules will greatly exceed EEEV RNA molecules particularly early after infection. The loss of a small number of miR-142-3p molecules due EEEV sequestration should not quantitatively change the levels of miR-142-3p within myeloid cells.

Our results also point to an effect of replication efficiency in initial infection sites on subsequent CNS disease severity. IFN-α/β production by the triple, quadruple and 11337 mutants at 12 hpi resulted in reduced virus dissemination to the CNS (Figure 5) suggesting that the initial innate immune responses within 12 hours after infection can influence events in tissues distant form the site of infection. We have previously shown production of serum IFN can lead to an upregulation of STAT1 transcription factor phosphorylation in the CNS [28]. Furthermore, WT EEEV and the 11337 mutant, missing all four sites, were equally virulent when virus was delivered ic. Therefore, the early IFN response in the triple, quadruple and 11337 deletion mutants may help to prime the innate immune response in the CNS to restrict virus replication and lessen disease severity. Myeloid cell production of cytokines and chemokines are also integral to the induction of both the innate and adaptive immune responses *in vivo* [26,27]. The inability of EEEV to replicate in tissue-specific myeloid cells may also lead to suppression of the cytokine and chemokine leading to inadequate induction of the adaptive immune responses in the spleen, or recruitment of important cell types to the CNS needed for neuroprotection. We infer that limitations in neuroprotective adaptive immune responses may also contribute to the extremely high mortality rate in symptomatic EEEV cases.

There is remarkable conservation of the EEEV 3’ UTRs that have been isolated from nature with a majority of the EEEV strains having nearly identical sequence and location of the miR-142-3p binding sites as the prototype strain EEEV FL93-939 [4]. Our data suggests that requirements for mosquito cell replication exert a strong selective pressure for maintenance of the miR-142-3p sites (current studies and [4]), which may explain such conservation. However, we found that in mammalian cells and hosts, the lack of a selective pressure results in generation of escape mutants that lack the miR-142-3p binding sites. Similarly, artificial miR-142-3p binding sites introduced into Dengue virus were deleted during the course of mouse infection [30]. The EEEV mutations we observed rendered all four of the miR-142-3p sites non-functional reinforcing the contribution of all sites to EEEV replication restriction.

Finally, we have shown that WEEV, a natural recombinant SINV/EEEV virus that derived its 3’ UTR sequences from an EEEV-like ancestor, contains four binding sites for miR-142-3p (Figure 6), but in different locations in the 3’ UTR compared to EEEV. Deletion of the binding sites from WEEV rescued virus replication in myeloid cells and attenuated the mutant virus in mice, while also suppressing replication in mosquito cells. Therefore, we suggest that multiple members of the encephalitic alphaviruses express virulence factors conferred by miRNA binding. It is of interest that the other major encephalitic alphavirus, Venezuelan equine encephalitis virus, has a much shorter 3’ UTR than EEEV or WEEV [45], does not possess miR-142-3p binding sites and is highly myeloid cell tropic [2].

In summary, our data demonstrate that the miR-142-3p binding sites cooperatively suppress EEEV 3’ UTR replication in myeloid cells thereby enhancing virulence *in vivo*. miRNA binding sites are being used to limit tissue tropism in the generation of live-attenuated vaccine candidates [22,23]. However, these artificial miRNA binding sites are usually complimentary to the entire miRNA [22,23,30], which is rarely found in naturally occurring viral RNAs [4,13–17]. Furthermore, when artificial miRNA binding sites are grouped together, viral escape of miRNA suppression can occur through deletion of the inserted miRNA binding sites [22,30], leading to potential adverse events.

## Materials and Methods

### Ethics Statement

All animal procedures were carried out under approval of the Institutional Animal Care and Use Committee of the University of Pittsburgh in protocols 15066059 and 18073259. Animal care and use were performed in accordance with the recommendations in the Guide for the Care and Use of Laboratory Animals of the National Research Council. Approved euthanasia criteria were based on weight loss and morbidity.

#### Cell culture

Baby hamster kidney cells (BHK-21), L929 fibroblasts, RAW 264.7 (RAW) monocyte-macrophage cells, and *Aedes albopictus* C6/36 mosquito cells were maintained as previously described[2,4]. Bone marrow-derived, conventional dendritic cells were generated from C57BL/6 mice (Jackson Laboratories) and maintained as previously described[2] in media supplemented with 10ng/ml recombinant interleukin-4 and 10ng/ml granulocyte-macrophage colony stimulating factor (Peprotech). Bone marrow macrophages (BMMΦ) were generated from *Ifnar*^−/−^ mice as previously described [2] in Dulbeccos’ modified Eagle’s medium supplemented with 20% L929-conditioned supernatant.

#### Viruses

Construction of the WT EEEV strain FL93-939 cDNA clone and mutant 11337 cDNA clone encoding nanoLuciferase (nLuc) as a cleavable in-frame fusion protein located between the capsid and E3 protein has been described previously [4,24,46]. EEEV mutant viruses containing three nucleotide mutations in the miR-142-3p binding sites in the 3’ UTR complimentary to the miR-142-3p seed sequence to eliminate miR-142-3p binding were generated singly or in combination using the QuikChange Mutagenesis II XL kit and the primers listed in S1 Table [4]. The infectious cDNA clone of WEEV McMillan strain (McM) was kindly provided by Kenneth Olson, Colorado State University[47]. This clone was modified by placing the entire virus sequence into the PBR-322 based vector of the FL93-939 virus cDNA under transcriptional control of the T7 bacteriophage promoter. A WEEV McM miR-142-3p mutant virus (WEEV 11224) was created by deleting 226 nucleotides in the 3’ UTR (nucleotide 11224 to 11449) by QuikChange Mutagenesis using the primers listed in S1 Table. Capped, *in vitro* transcribed RNA was generated from the linearized cDNA using the T7 mMessage mMachine kit (Ambion), and electroporated into BHK cells as previously described [2]. Titers of virus stocks was determined by BHK-21 cell plaque assay.

#### Translation Reporters and RNA immunoprecipitations

Generation of the WT EEEV, and 11337 translation reporters were described previously [4]. Capped, *in vitro* transcribed RNA was generated for use in immunoprecipitation assays. Raw cells were transfected with 7ug reporter RNAs either with or without 30 pmoles of biotinylated miR-142-3p mimic or scrambled 3’ biotinylated mimic (Dharmacon) using the Neon Transfection System (Invitrogen; 1750V, 25ms, 1 pulse). At 1.5-2h post-transfection, cells were washed thrice using ice-cold PBS w/o Ca^+^ and Mg^+^. Lysates were collected on ice in modified radioimmunoprecipitation assay (RIPA) buffer (50 mM Tris, pH 7.4, 150 mM NaCl, 1% NP-40) supplemented with protease inhibitors (1 mM phenylmethylsulfonyl fluoride, 1μg/ml leupeptin, and 1 μg/ml pepstatin), and a phosphatase inhibitor cocktail (Sigma). Lysates were spun at 12000g for 10 min at 4°C to clear debris and supernatants transferred to pre-chilled tubes. For immunoprecipitation of the Argonaute complex, Protein A/G Plus agarose beads were blocked with bovine t-RNA (1mg/mL; Sigma) for 2h in modified-RIPA buffer, washed 2X and resuspended in modified-RIPA buffer. Lysates were pre-cleared by adding 1μL rabbit serum and 30μL Protein A/G Plus agarose beads (30 μL per sample; Santa Cruz) for 2h at 4°C on a nutator. Lysates were spun at 2500g for 10 min at 4°C and supernatants transferred to pre-chilled tubes. Anti-eiF2C rabbit polyclonal antibody (30 μL per sample; Santa Cruz; H-300) was added and rocked on a nutator O/N at 4°C. t-RNA blocked protein A/G beads were added for an additional 2h at 4°C. Lysates were spun at 2500g and washed 3X in ice-cold RIPA buffer. Beads were suspended in Trizol reagent (Ambion) and freeze-thawed at −80°C. For immunoprecipitation of the biotinylated mimic RNA, streptavidin agarose beads (30μL per sample; Cell Signaling) were blocked with bovine t-RNA (1mg/mL; Sigma) for 2h in RIPA buffer, washed 2X and resuspended in RIPA buffer. t-RNA blocked streptavidin beads were added for 6h at 4°C on a nutator. Lysates were spun at 2500g and washed 3X in ice cold RIPA buffer. Beads were suspended in Trizol reagent (Ambion) and freeze-thawed at −80°C. For both immunoprecipitations, RNA was extracted using manufacturers guidelines (Ambion). 100 ng of total RNA was used in a reverse transcription reaction with random hexamer (Integrated DNA Technologies), and resultant cDNA was used to detect levels of reporter RNAs with the following primers: sense (5’-GGGAGCGCGCCTGTAAGGCACAC-3’) and antisense (5’-GCTCTCCAGCGGTTCCATCTTCCAGC-3’). Data was normalized to mock samples and total input RNA levels.

#### Virus Growth Curves

BHK (2×10^5^ cells/well), RAW (2×10^5^ cells/well), BMDCs (1.5×10^5^ cells/well), and C6/36 cells (2×10^5^ cells/well) were seeded in 24 well plates one day prior to infection. Viruses were infected in triplicate at a MOI = 0.1 for RAW and BHK cells, MOI = 5 for BMDCs, or MOI = 1 for C6/36 cells in phosphate buffered saline (PBS) supplemented with 1% FBS. After 1 hour, the cells were washed with PBS and complete media was added to each well. For nLuc analysis, at the indicated time points, the cells were washed three times with PBS followed by addition of 100ul of 1X Passive lysis buffer (PLB, Promega). The cells were then scraped and transferred to a 96 well plate. For plaque assay, supernatant was collected at time zero and indicated time points for titration by plaque assay on BHK-21 cells.

#### Mouse infection and tissue collection

6-week old female outbred CD-1 mice (Charles River Laboratories) or C57BL/6J (Jackson Laboratories) were infected subcutaneously (sc) in each footpad with 10^3^ pfu of nLuc-expressing EEEV mutant viruses in 10 ul of OptiMEM media (Invitrogen). For WEEV, female *Ifnar*^−/−^ and CD-1 mice were infected with 10^4^ pfu of WEEV McM or WEEV 11224 sc in each footpad. All mice were scored daily for clinical signs and weight loss as described previously [2]. For tissue collection, serum was collected via the submandibular vein, and the mice were perfused with PBS. Tissues were harvested and collected into Eppendorf tubes containing 1x PLB (e.g.100ul per a single popliteal lymph node (PLN), 400ul per footpad, 800ul per spleen and brain), or Tri-Reagent for RNA analysis. For aerosol infection, CD-1 mice were challenged with 100LD_50_ of EEEV FL93 as previously described [25]. On day 5 post infection, cervical lymph nodes (CVLN) were harvested and processed in Tri-Reagent prior to being frozen at −80°C. Samples in PLB were homogenized and refrozen at −80°C prior to analysis. All animal procedures were carried out in accordance with the American Association for the Accreditation of Laboratory Animal Care International-approved institutional guidelines for animal use and approved by the University of Pittsburgh Institutional Animal Care and Use Committee.

### Luciferase Assays and Protein Assays

nLuc assays were using the Nano-Glo Luciferase system (Promega) and preformed according to manufacturer’s guidelines. Samples were diluted in 1x PLB for determination of nLuc relative light units (RLU) using a Orion microplate luminometer (Berthold) or a FLUOstar Omega microplate reader (BMG Labtech). RLU was normalized to protein levels in samples determined by a bicinchoninic acid protein assay (Pierce).

### Interferon (IFN- α/β) Bioassays

Biologically active serum IFN-α/β collected at 12 and 24 hpi was measured using a standard IFN biological assay on L929 cells as described previously [48]. The IFN-α/β concentration in sera samples was set as the dilution of sample required for 50% protection from cytopathic effect compared to protection conferred by an IFN standard [25].

### RNA Isolation and RT-PCR

RNA was isolated from PLN in Tri-reagent according to manufacturer’s guidelines. Poly-acryl carrier was added to each PLN prior to addition of 1-bromo-3-choropropane (BCP) and phase separation. Reverse transcription (RT) of 100 ng of RNA was performed as previously described [49] using Moloney Murine Leukemia virus (M-MLV) reverse transcriptase (Promega), with an extension temperature of 42°C for 60 min and random hexamer (IDT). For cytokine and chemokine analysis, Maxima qPCR SYBR Green/ROX Master Mix (ThermoFisher) was used and the primers in S2 Table. Threshold cycle (*C*_*T*_) values were normalized to 18s and compared to mock samples using the ΔΔ*C*_*T*_ method.

### Identification of EEEV Escape Mutants

RNA was isolated as described from *in vitro* cultured RAW cells or BMDCs at 48 hr post infection or EEEV-infected (1×10^3^ pfu bilaterally in footpad) and brain (D5), serum (24 hpi), and CVLN (D5) samples were harvested and placed in Tri-Reagent for RNA analysis. RT was performed using 50uM Oligo(dT) (Thermo Fisher) as previously described [49] with an 48°C extension temperature. cDNA was diluted with H_2_O and 10ul was used in a GoTaq PCR reaction (GoTaq Green Mastermix, Fisher Scientific) with the following conditions: 95°C 2 min, (95°C 45s, 60°C 30s, 73°C 60s) x 40 cycles, 73°C 7 min. The following primers were used: EEEV 10951-S: CGTTGCCTACAAATCCAGTAAAGCAGGA; T7-EEECSE-AS: TAATACGACTCACTATAGGGCGTATGGAAAAAATTAATATGATTTTGTAAATTGATATAAAAGACAGC. The entire PCR reaction was run on a 2% agarose TAE gel followed by excising the bands and clean-up using Wizard SV Gel Clean-up (Promega). cDNA from WT EEEV and 11337 stocks were used as positive controls during PCR. Sequencing was performed by the University of Pittsburgh HSCRF Genomics Research Core and analyzed using CLC Genomics Workbench (Qiagen).

### Software and Statistical Analysis

miRANDA-3.3a software [50,51] was used to align the mmu-miR-142-3p sequence with the EEEV FL93 genome (EF151502.1) and WEEV McMillan genome (GQ287640.1). All statistical analysis was performed using GraphPad Prism software. All experiments were repeated at least twice as indicated in Figure Legends. For IP, unpaired t test was performed to compared between groups. For the EEEV point mutant *in vitro* and *in vivo* data, a one-way analysis of variance was performed of the log-transformed data with corrections for multiple comparison using the Holm-Sidak method comparing each mutant to WT. An unpaired t-test was used to compare 11337 to mutant 1234 in the growth curve experiments in C6/36 cells. Box-and whisker plots represent min-max with bar representing the median value. Statistical significance for survival curves was determined by Mantel-Cox log rank test compared to WT. For the WEEV growth curves, multiple unpaired t tests were perfumed and corrected for multiple comparisons using the Holm-Sidak method.

## Acknowledgements

We would like to thank Chelsea Maksin for her excellent technical assistance.

## Supporting Information Captions

**S1 Figure. EEEV mutants lacking functional miR-142-3p binding sites are attenuated in C57BL6 mice.** Female C57BL6 mice (5-6 weeks) were infected with 10^3^ pfu sc in each footpad. Morbidity and mortality were measured twice daily. n=7-8 mice from 2 independent experiments

**S2 Figure: Increased myeloid cell replication leads to increased cytokine and chemokine mRNA in PLN.** A) Cytokine and chemokine mRNA levels in the PLN of CD-1 mice 12 hpi with 10^3^ pfu of WT, 11337 or mock infected. Data is represented as fold difference compared to mock mice. n=12 mice, 3 independent experiments B) *Cxcl10* mRNA levels in PLN 12 hpi with EEEV mutants. n=8-12 mice, 2-3 independent experiments *P<0.05, **P<0.01, ***P<0.001, ****P<0.0001, (A) one way analysis of variance test with corrections for multiple comparisons using Turkey method or (B) one way analysis of variance test between WT and each mutant with corrections for multiple comparisons using Holm-Sidak method of the log-transformed data. ns=non-significant. Box-and whisker plots represent min-max with bar representing the median value.

**S3 Figure: Reduced virus replication in the periphery with the triple and quadruple mutant EEEV viruses.** CD-1 mice were infected with 10^3^ pfu of the EEEV mutants sc in each footpad. Tissues were harvested at 96 hours post infection. Virus replication in PLN (A), spleen (B). N=8 mice, from 2 independent experiments. *P<0.5, **P<0.01, ***P<0.001, ****P<0.0001 one way analysis of variance test with corrections for multiple comparisons using the Holm-Sidak method comparing each mutant to WT. Box-and whisker plots represent min-max with bar representing the median value.

**S4 Figure: Mutant 11337 is virulent after intracerebral infection.** Survival of female (5-6 week) CD-1 infected with ic with either 10^3^ pfu of WT or 11337 mutant. Morbidity and mortality were measured twice daily. n=8 mice from 2 independent experiments.

**S5 Figure: Escape mutants generated during infection eliminate miR-142-3p binding sites in EEEV 3’ UTR.** Alignment of escape mutants isolated for indicated cells or tissues. Numbers on left indicate location in the genome of the deletion. Numbers at end of the sequence indicate the length of the 3’ UTR in each escape mutant. B6 - C57Bl6, BMDC- bone marrow derived dendritic cell, CVLN – cervical lymph nodes

## References

1. Deresiewicz RL, Thaler SJ, Hsu L, Zamani AA. Clinical and neuroradiographic manifestations of eastern equine encephalitis. N Engl J Med. 1997;336: 1867–1874. doi:10.1056/NEJM199706263362604

2. Gardner CL, Burke CW, Tesfay MZ, Glass PJ, Klimstra WB, Ryman KD. Eastern and Venezuelan equine encephalitis viruses differ in their ability to infect dendritic cells and macrophages: impact of altered cell tropism on pathogenesis. J Virol. 2008;82: 10634–10646. doi:10.1128/JVI.01323-08

3. Mildner A, Chapnik E, Manor O, Yona S, Kim K-W, Aychek T, et al. Mononuclear phagocyte miRNome analysis identifies miR-142 as critical regulator of murine dendritic cell homeostasis. Blood. 2013;121: 1016–1027. doi:10.1182/blood-2012-07-445999

4. Trobaugh DW, Gardner CL, Sun C, Haddow AD, Wang E, Chapnik E, et al. RNA viruses can hijack vertebrate microRNAs to suppress innate immunity. Nature. 2014;506: 245–248. doi:10.1038/nature12869

5. Bartel DP. MicroRNAs: genomics, biogenesis, mechanism, and function. Cell. 2004;116: 281–297.

6. Hutvágner G, Zamore PD. A microRNA in a multiple-turnover RNAi enzyme complex. Science. American Association for the Advancement of Science; 2002;297: 2056–2060. doi:10.1126/science.1073827

7. Zeng Y, Yi R, Cullen BR. MicroRNAs and small interfering RNAs can inhibit mRNA expression by similar mechanisms. Proc Natl Acad Sci USA. 2003;100: 9779–9784. doi:10.1073/pnas.1630797100

8. Bartel DP. MicroRNAs: Target Recognition and Regulatory Functions. Cell. 2009;136: 215–233. doi:10.1016/j.cell.2009.01.002

9. Agarwal V, Bell GW, Nam J-W, Bartel DP. Predicting effective microRNA target sites in mammalian mRNAs. Elife. 2015;4: 101. doi:10.7554/eLife.05005

10. Guo H, Ingolia NT, Weissman JS, Bartel DP. Mammalian microRNAs predominantly act to decrease target mRNA levels. Nature. 2010;466: 835–840. doi:10.1038/nature09267

11. Eichhorn SW, Guo H, McGeary SE, Rodriguez-Mias RA, Shin C, Baek D, et al. mRNA destabilization is the dominant effect of mammalian microRNAs by the time substantial repression ensues. Mol Cell. 2014;56: 104–115. doi:10.1016/j.molcel.2014.08.028

12. Jopling CL, Yi M, Lancaster AM, Lemon SM, Sarnow P. Modulation of hepatitis C virus RNA abundance by a liver-specific MicroRNA. Science. 2005;309: 1577–1581. doi:10.1126/science.1113329

13. Scheel TKH, Luna JM, Liniger M, Nishiuchi E, Rozen-Gagnon K, Shlomai A, et al. A Broad RNA Virus Survey Reveals Both miRNA Dependence and Functional Sequestration. Cell Host Microbe. 2016;19: 409–423. doi:10.1016/j.chom.2016.02.007

14. Song L, Liu H, Gao S, Jiang W, Huang W. Cellular microRNAs inhibit replication of the H1N1 influenza A virus in infected cells. J Virol. 2010;84: 8849–8860. doi:10.1128/JVI.00456-10

15. Khongnomnan K, Makkoch J, Poomipak W, Poovorawan Y, Payungporn S. Human miR-3145 inhibits influenza A viruses replication by targeting and silencing viral PB1 gene. Exp Biol Med (Maywood). SAGE Publications; 2015;240: 1630–1639. doi:10.1177/1535370215589051

16. Ingle H, Kumar S, Raut AA, Mishra A, Kulkarni DD, Kameyama T, et al. The microRNA miR-485 targets host and influenza virus transcripts to regulate antiviral immunity and restrict viral replication. Sci Signal. 2015;8: ra126. doi:10.1126/scisignal.aab3183

17. Zheng Z, Ke X, Wang M, He S, Li Q, Zheng C, et al. Human microRNA hsa-miR-296-5p suppresses enterovirus 71 replication by targeting the viral genome. J Virol. 2013;87: 5645–5656. doi:10.1128/JVI.02655-12

18. Wen B-P, Dai H-J, Yang Y-H, Zhuang Y, Sheng R. MicroRNA-23b Inhibits Enterovirus 71 Replication through Downregulation of EV71 VPl Protein. Intervirology. 2013;56: 195–200. doi:10.1159/000348504

19. Trobaugh DW, Klimstra WB. MicroRNA Regulation of RNA Virus Replication and Pathogenesis. Trends Mol Med. 2017;23: 80–93. doi:10.1016/j.molmed.2016.11.003

20. Shimakami T, Yamane D, Jangra RK, Kempf BJ, Spaniel C, Barton DJ, et al. Stabilization of hepatitis C virus RNA by an Ago2-miR-122 complex. Proc Natl Acad Sci USA. 2012;109: 941–946. doi:10.1073/pnas.1112263109

21. Meister G. Argonaute proteins: functional insights and emerging roles. Nat Rev Genet. Nature Publishing Group; 2013;14: 447–459. doi:10.1038/nrg3462

22. Heiss BL, Maximova OA, Thach DC, Speicher JM, Pletnev AG. MicroRNA Targeting of Neurotropic Flavivirus: Effective Control of Virus Escape and Reversion to Neurovirulent Phenotype. J Virol. 2012;86: 5647–5659. doi:10.1128/JVI.07125-11

23. Teterina NL, Liu G, Maximova OA, Pletnev AG. Silencing of neurotropic flavivirus replication in the central nervous system by combining multiple microRNA target insertions in two distinct viral genome regions. Virology. 2014;456-457: 247–258. doi:10.1016/j.virol.2014.04.001

24. Sun C, Gardner CL, Watson AM, Ryman KD, Klimstra WB. Stable, high-level expression of reporter proteins from improved alphavirus expression vectors to track replication and dissemination during encephalitic and arthritogenic disease. J Virol. 2014;88: 2035–2046. doi:10.1128/JVI.02990-13

25. Trobaugh DW, Sun C, Dunn MD, Reed DS, Klimstra WB. Rational design of a live-attenuated eastern equine encephalitis virus vaccine through informed mutation of virulence determinants. PLoS Pathog. 2019;15: e1007584. doi:10.1371/journal.ppat.1007584

26. Shi C, Pamer EG. Monocyte recruitment during infection and inflammation. Nat Rev Immunol. 2011;11: 762–774. doi:10.1038/nri3070

27. Jain A, Pasare C. Innate Control of Adaptive Immunity: Beyond the Three-Signal Paradigm. J Immunol. American Association of Immunologists; 2017;198: 3791–3800. doi:10.4049/jimmunol.1602000

28. Gardner CL, Yin J, Burke CW, Klimstra WB, Ryman KD. Type I interferon induction is correlated with attenuation of a South American eastern equine encephalitis virus strain in mice. Virology. Elsevier Inc; 2009;390: 338–347. doi:10.1016/j.virol.2009.05.030

29. Armstrong PM, Andreadis TG. Eastern Equine Encephalitis Virus in Mosquitoes and Their Role as Bridge Vectors. Emerg Infect Dis. 2010;16: 1869–1874. doi:10.3201/eid1612.100640

30. Pham AM, Langlois RA, tenOever BR. Replication in Cells of Hematopoietic Origin Is Necessary for Dengue Virus Dissemination. Kuhn RJ, editor. PLoS Pathog. 2012;8: e1002465. doi:10.1371/journal.ppat.1002465.s005

31. Yao Y, Charlesworth J, Nair V, Watson M. MicroRNA expression profiles in avian haemopoietic cells. Front Genet. Frontiers; 2013;4: 153. doi:10.3389/fgene.2013.00153

32. Hahn CS, Lustig S, Strauss EG, Strauss JH. Western equine encephalitis virus is a recombinant virus. Proc Natl Acad Sci USA. 1988;85: 5997–6001.

33. Weaver SC, Kang W, Shirako Y, Rumenapf T, Strauss EG, Strauss JH. Recombinational history and molecular evolution of western equine encephalomyelitis complex alphaviruses. J Virol. American Society for Microbiology (ASM); 1997;71: 613–623.

34. Levitt NH, Miller HV, Edelman R. Interaction of alphaviruses with human peripheral leukocytes: in vitro replication of Venezuelan equine encephalomyelitis virus in monocyte cultures. Infect Immun. American Society for Microbiology (ASM); 1979;24: 642–646.

35. Langlois RA, Varble A, Chua MA, García-Sastre A, tenOever BR. Hematopoietic-specific targeting of influenza A virus reveals replication requirements for induction of antiviral immune responses. Proc Nat Acad Sci. National Acad Sciences; 2012;109: 12117–12122. doi:10.1073/pnas.1206039109

36. Langlois RA, Albrecht RA, Kimble B, Sutton T, Shapiro JS, Finch C, et al. MicrorNA-based strategy to mitigate the risk of gain-of-function influenza studies. Nat Biotechnol. Nature Publishing Group; 2013;31: 844–847. doi:10.1038/nbt.2666

37. Hon LS, Zhang Z. The roles of binding site arrangement and combinatorial targeting in microRNA repression of gene expression. Genome Biol. BioMed Central; 2007;8: R166. doi:10.1186/gb-2007-8-8-r166

38. Grimson A, Farh KK-H, Johnston WK, Garrett-Engele P, Lim LP, Bartel DP. MicroRNA targeting specificity in mammals: determinants beyond seed pairing. Mol Cell. 2007;27: 91–105. doi:10.1016/j.molcel.2007.06.017

39. Krek A, Grün D, Poy MN, Wolf R, Rosenberg L, Epstein EJ, et al. Combinatorial microRNA target predictions. Nat Genet. 2005;37: 495–500. doi:10.1038/ng1536

40. Saetrom P, Heale BSE, Snøve O, Aagaard L, Alluin J, Rossi JJ. Distance constraints between microRNA target sites dictate efficacy and cooperativity. Nucleic Acids Res. 2007;35: 2333–2342. doi:10.1093/nar/gkm133

41. Lecellier C-H, Dunoyer P, Arar K, Lehmann-Che J, Eyquem S, Himber C, et al. A cellular microRNA mediates antiviral defense in human cells. Science. 2005;308: 557–560. doi:10.1126/science.1108784

42. Bai XT, Nicot C. miR-28-3p Is a Cellular Restriction Factor That Inhibits Human T Cell Leukemia Virus, Type 1 (HTLV-1) Replication and Virus Infection. J Biol Chem. 2015;290: 5381–5390. doi:10.1074/jbc.M114.626325

43. Friedman RC, Farh KK-H, Burge CB, Bartel DP. Most mammalian mRNAs are conserved targets of microRNAs. Genome Res. 2009;19: 92–105. doi:10.1101/gr.082701.108

44. Luna JM, Scheel TKH, Danino T, Shaw KS, Mele A, Fak JJ, et al. Hepatitis C Virus RNA Functionally Sequesters miR-122. Cell. 2015;160: 1099–1110. doi:10.1016/j.cell.2015.02.025

45. Hyde JL, Chen R, Trobaugh DW, Diamond MS, Weaver SC, Klimstra WB, et al. The 5“ and 3” ends of alphavirus RNAs--Non-coding is not non-functional. Virus Res. 2015;206: 99–107. doi:10.1016/j.virusres.2015.01.016

46. Aguilar PV, Adams AP, Wang E, Kang W, Carrara A-S, Anishchenko M, et al. Structural and nonstructural protein genome regions of eastern equine encephalitis virus are determinants of interferon sensitivity and murine virulence. J Virol. 2008;82: 4920–4930. doi:10.1128/JVI.02514-07

47. Logue CH, Bosio CF, Welte T, Keene KM, Ledermann JP, Phillips A, et al. Virulence variation among isolates of western equine encephalitis virus in an outbred mouse model. J Gen Virol. 2009;90: 1848–1858. doi:10.1099/vir.0.008656-0

48. Bhalla N, Sun C, Metthew Lam LK, Gardner CL, Ryman KD, Klimstra WB. Host translation shutoff mediated by non-structural protein 2 is a critical factor in the antiviral state resistance of Venezuelan equine encephalitis virus. Virology. 2016;496: 147–165. doi:10.1016/j.virol.2016.06.005

49. Watson AM, Lam LKM, Klimstra WB, Ryman KD. The 17D-204 Vaccine Strain-Induced Protection against Virulent Yellow Fever Virus Is Mediated by Humoral Immunity and CD4+ but not CD8+ T Cells. Pierson TC, editor. PLoS Pathog. 2016;12: e1005786–29. doi:10.1371/journal.ppat.1005786

50. Enright AJ, John B, Gaul U, Tuschl T, Sander C, Marks DS. MicroRNA targets in Drosophila. Genome Biol. 2003;5: R1. doi:10.1186/gb-2003-5-1-r1

51. John B, Enright AJ, Aravin A, Tuschl T, Sander C, Marks DS. Human MicroRNA targets. James C Carrington, editor. Plos Biol. Public Library of Science; 2004;2: e363. doi:10.1371/journal.pbio.0020363

